## Supplementary figures and images for "Caveolin-1 protects endothelial cells from extensive expansion of transcellular tunnel by stiffening the plasma membrane"

### Supplementary Figures

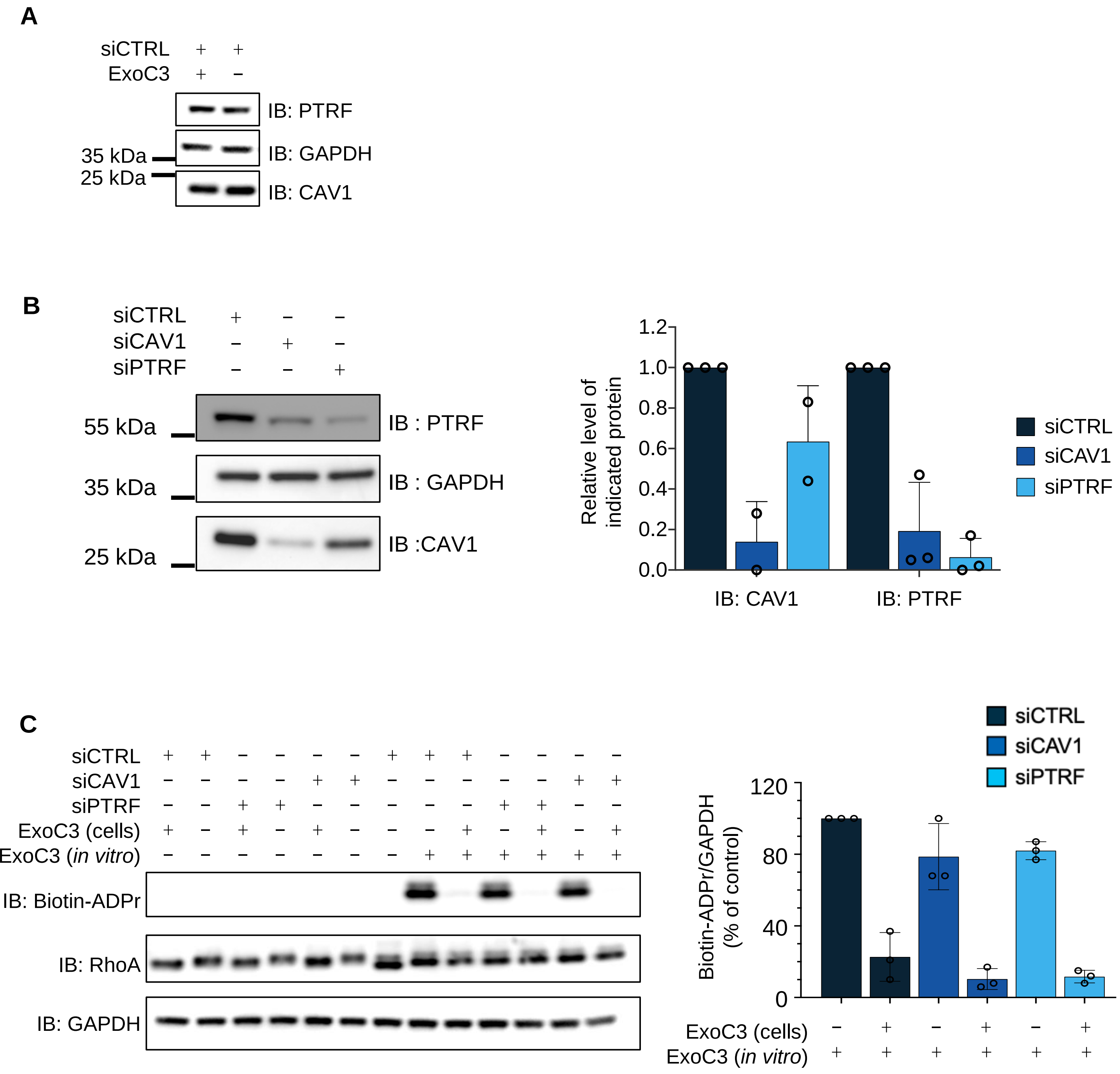

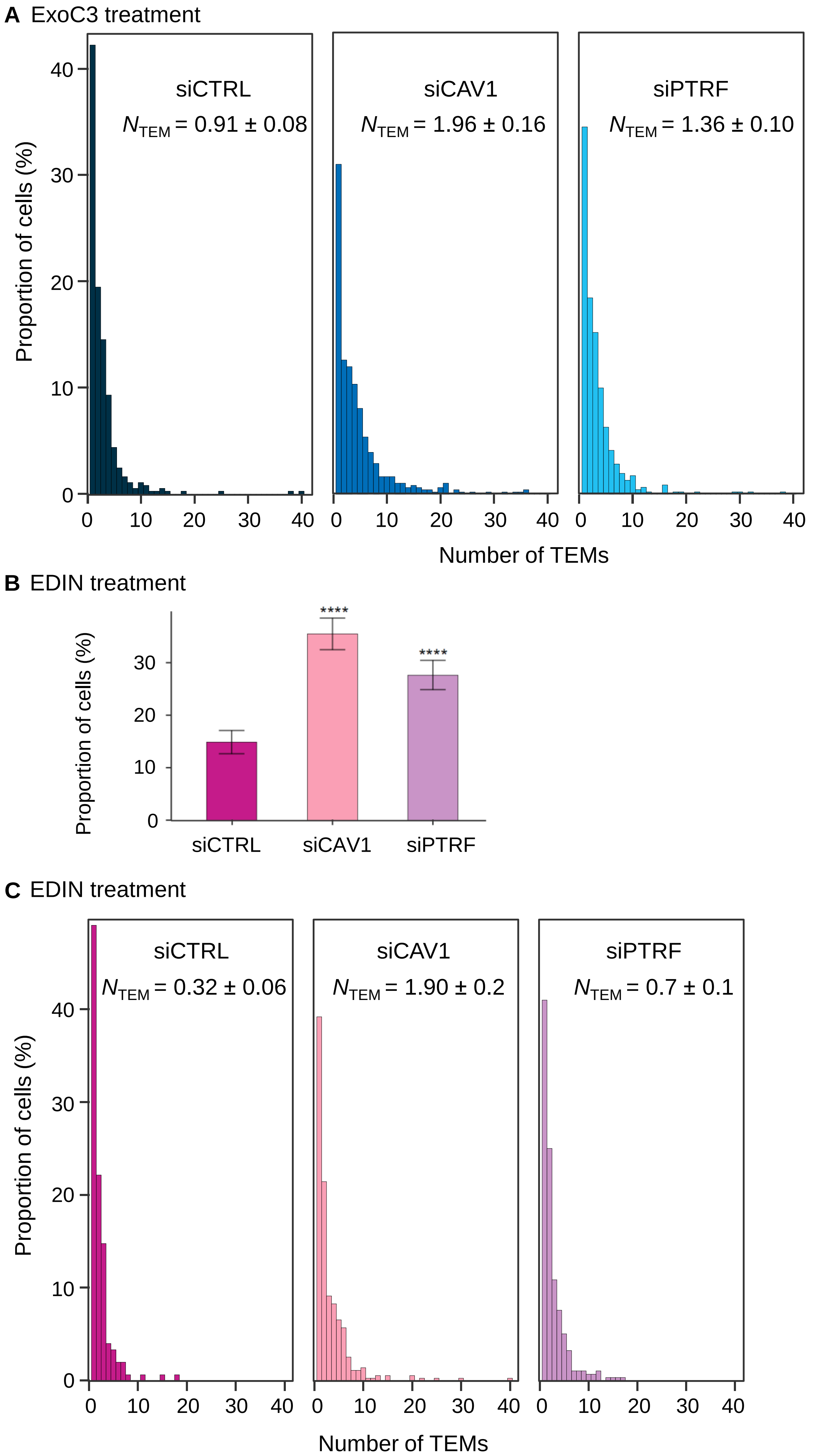

Figure S3

A

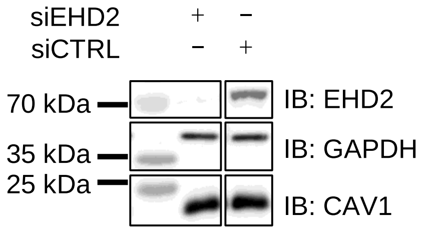


B ExoC3 treatment

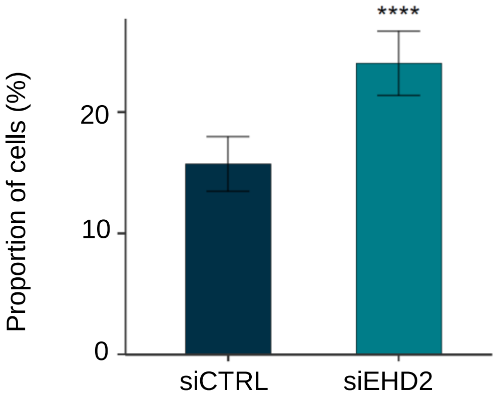


C ExoC3 treatment

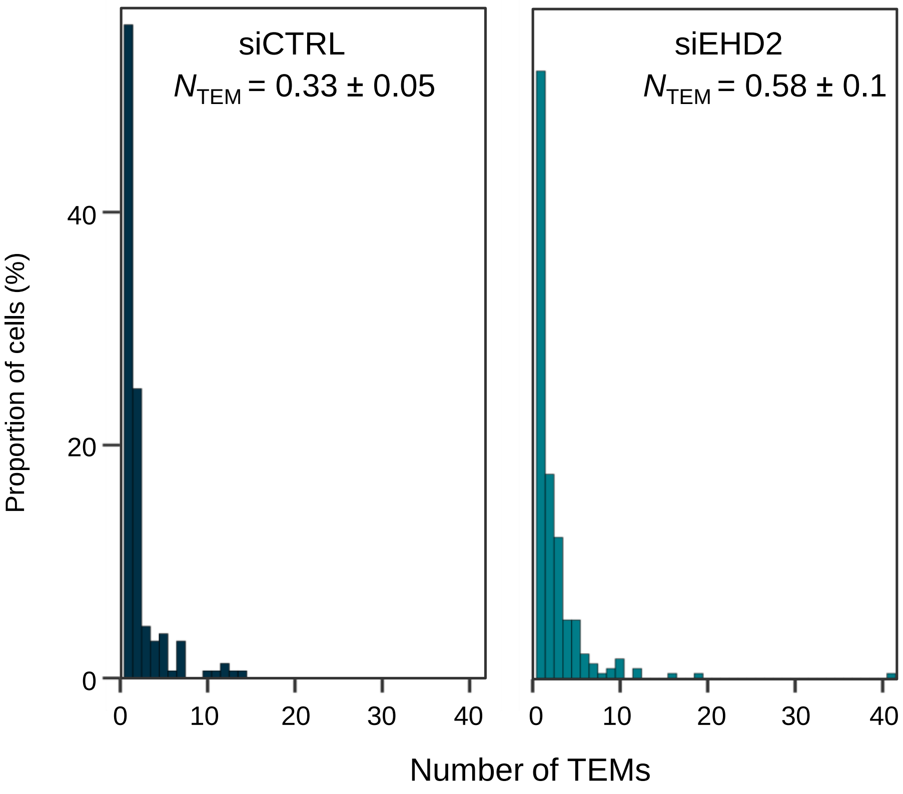

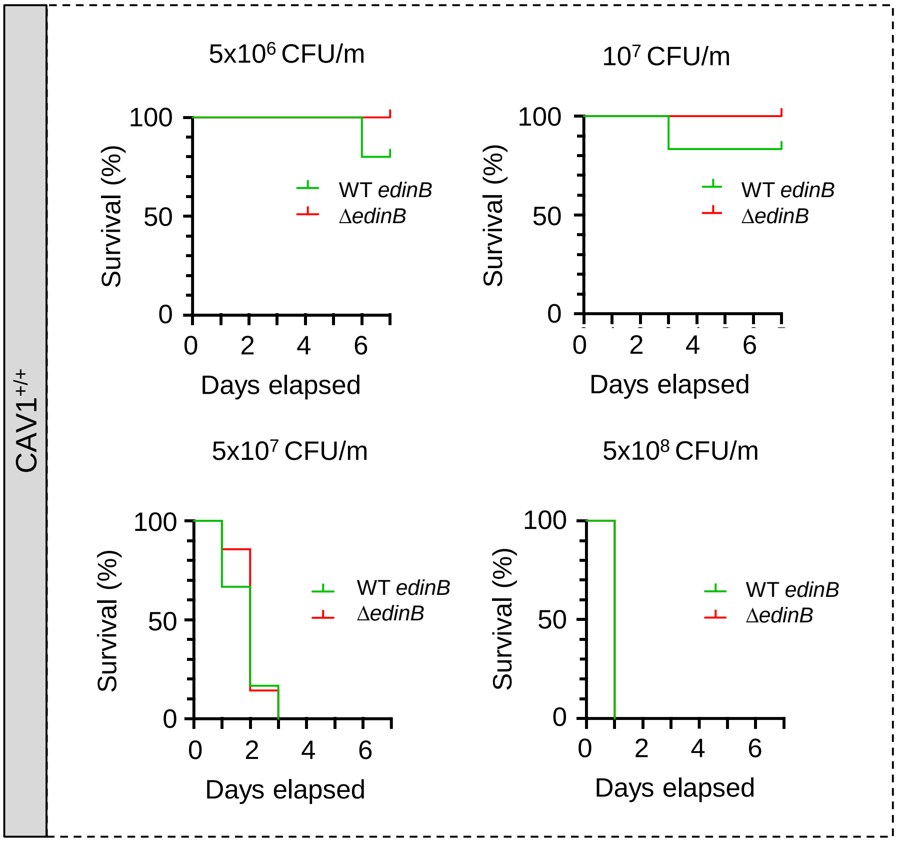

Figure S5

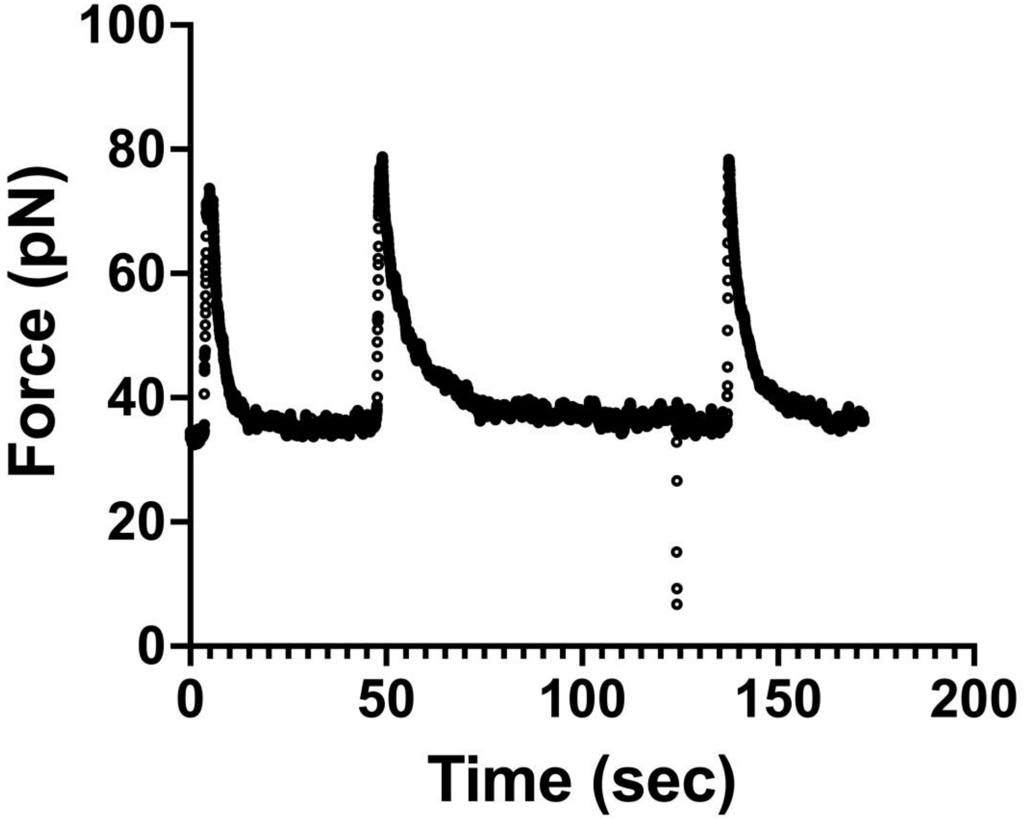
